# Do urban ecosystem service assessments account for ecosystem condition and biodiversity?

**DOI:** 10.64898/2025.12.03.692179

**Authors:** Davide Stucchi, Renato Casagrandi, Javier Babí Almenar

## Abstract

Urban ecosystems provide essential ecosystem services (ES), a supply that is dependent on ecosystem condition (EC). Statistical frameworks like the UN SEEA-EA explicitly link ES flows to the extent and condition of ecosystem assets. Yet, the concrete role of EC in shaping ES supply remains only partially understood, as their relationship is complex and varies across services. Consequently, the operational integration of EC into urban ES assessments remains fragmented. Through a systematic review of 110 studies (2005-2024), we evaluated how and to what extent EC has been incorporated into urban ES assessments. For each study, we examined ES assessment methods, types of condition variables, spatial and temporal explicitness, flow types (potential vs. actual), and sustainability considerations. We find that integration is decisively underway, with most studies (87%) using EC variables as inputs, predominantly for regulating services. However, this integration is narrow: it relies on static methods, focuses on potential over actual flows, and favors easily measurable abiotic and structural state variables over functional, compositional, or landscape ones. While spatial explicitness is common (45%), dynamic models are rare (20%), and assessments seldom leverage EC to evaluate ES flow sustainability. Addressing these gaps, by broadening types of EC variables, increasing temporal dynamism, and linking condition to both actual flows and sustainability, will enhance the capacity of urban ES assessments to support the planning and adaptive management of urban greening. It will also help advance the development of urban ecosystem condition and service accounts, thereby increasing the relevance of urban ES assessment knowledge.

## 1. Introduction

Urban ecosystems are complex socio-ecological systems that combine human presence with a fine grained mosaic of natural, semi-natural, and artificial components (Grimm et al., 2008; United Nations, 2021; Wu, 2014). They are increasingly recognized as critical providers of ecosystem services (ES), as reflected in the rapid expansion of urban ES research (Shao et al., 2023), and the growing promotion of concepts like green infrastructure, nature-based solutions, and ecosystem assets by policymakers, researchers, and practitioners (Frantzeskaki et al., 2020; Grace et al., 2021; Keeler et al., 2019; Venter et al., 2020). These concepts typically frame urban greening and its components (e.g., trees, shrubs, ponds, bioswales) as features, solutions, or assets that generate ES supply, thereby contributing to generate human benefits and helping mitigate urban challenges such as urban heat island or poor air quality, strongly shaped by abiotic and biotic characteristics (Babí Almenar et al., 2021; Eggermont et al., 2015). Thus, effective planning and management of urban greening requires quantifying their supply over time.

Quantifying ES necessitates understanding the characteristics and factors that shape ES supply– demand dynamics and how they evolve over time (Babí Almenar et al., 2023). Central to this is ecosystem condition, as a key driver of ES supply (La Notte et al., 2019; Sutherland et al., 2018). Ecosystem condition (EC) represents the quality of an ecosystem in terms of intrinsic, relational, and instrumental values, measured through abiotic and biotic characteristics across a range of temporal and spatial scales (Czúcz et al., 2021; Keith et al., 2020). From an instrumental perspective, EC partly determines the capacity of an ecosystem asset to generate ES, as changes in condition (e.g., degradation) are expected to affect ES flows (Czúcz et al., 2021; European Commission et al., 2014; United Nations, 2021). Although, EC-ES relationships are often complex, non-linear or delayed, and vary across services, ecosystems, and contexts, the dependency of ES supply on EC is firmly established in frameworks such as the ecosystem cascade (La Notte et al., 2017). Likewise, the UN System of Environmental-Economic Accounting—Ecosystem Accounting (SEEA-EA) conceptualizes ES supply flows as dependent on the extent and condition of ecosystem assets (United Nations, 2021), with urban greening being such assets within urban ecosystems.

Despite this conceptual clarity, the operational integration of EC into urban ES assessments remains fragmented. Haase et al., (2014) observed that most studies rely on land use/land cover (LULC) changes as proxies for ES supply. For instance, Hamel et al., (2021) used InVEST urban tools to map multiple ES types, but the differences in outputs across scenarios reflected mostly LULC changes. Widely used tools such as InVEST, ARIES, and LUCI rely predominantly on LULC inputs, which insufficiently capture the condition-dependent nuances of many regulating services (Cabral et al., 2016; Martínez-López et al., 2019; Peh et al., 2013; Trodahl et al., 2017). More advanced models, such as i-Tree, ENVI-met, and SWMM, incorporate biophysical variables (e.g., leaf area index, soil texture) and can better detect changes in ES supply over time (Burszta-Adamiak and Mrowiec, 2013; Hirabayashi et al., 2015; Zölch et al., 2016). However, they often treat EC variables as static factors, missing their temporal dynamics. Furthermore, many urban ES assessment link condition variables directly to individual ES rather than acknowledging their influence across multiple services or capturing ES supply–demand dynamics (Ouyang and Luo, 2022). As a result, even when EC variables are included, it remains unclear whether existing methods sufficiently account for spatio-temporal variability or its implications for synergies and trade-offs in ES supply.

While there are previous reviews that have examined factors influencing ES supply more broadly (Bordt and Saner, 2019; Smith et al., 2017) or variables used to assess and map condition across ecosystem types (Nicholson Thomas et al., 2025), to our knowledge no recent systematic review has focused specifically on how condition variables are considered in urban ES assessments. A deeper understanding of the role of EC in urban ES supply is therefore needed. Such knowledge would help identify which characteristics of urban greening most strongly affect ES supply and ecosystem capacity. It is also essential to support an effective planning and management of urban greening, and entire urban ecosystems, to deliver a suitable range of ES while avoiding overexploitation and minimizing unmet demand (La Notte, 2024; La Notte et al., 2019).

A clear understanding of EC is also critical for developing robust urban ecosystem accounts under SEEA-EA, which require spatially explicit account for condition variables and information on how changes in EC influence ecosystem capacity and ES supply over time. Urban ecosystems are particularly relevant from an ecosystem accounting perspective: as primary human habitats and hotspots of ES demand, they are major drivers of global and local biodiversity change (Elliot et al., 2022b; IPBES et al., 2019; Isbell et al., 2023; Maxwell et al., 2016). Yet, despite recent works (European Commission. Joint Research Centre., 2022; Nicholson Thomas et al., 2025), it remains unclear which EC variables are most suitable for monitoring urban ecosystems and tracking their changes.

To address these gaps, this paper conducts a systematic review of the role of EC in urban ES assessments. Here, EC is inclusively defined to encompass biodiversity variables, whether or not they are conventionally treated as condition metrics. The overarching research question guiding the review is: *How, and to what extent, do urban ES assessments consider EC?* This question is unpacked into five specific questions, addressing the *how* (RQ 1-3) and the *to what extent* (RQ 4-5):

RQ1. Is EC integrated into urban ES assessment methods, or are both assessments merely coupled?
RQ2. Do urban ES assessments that consider EC estimate potential or actual flows, and do they consider the sustainability of the flows?
RQ3. When EC is integrated, are urban ES assessment methods spatially explicit and/or temporally dynamic?
RQ4. For which ES classes and types of methods is EC considered?
RQ5. Which types of EC variables are included in urban ES assessments?

Answering these questions will help clarify the current state-of-the-art in integrating EC into urban ES studies, identify critical gaps, and provide relevant insights for advancing the modelling of urban greening and its ES supply, as well as implementing SEEA-EA urban ecosystem accounts.

## 2. Methods

The systematic literature review was designed to identify and select studies that assess ES in urban contexts and that either couple or integrate EC variables within such assessments. In this review, *coupled* refers to studies that assess ES and EC independently but in relation to one another, whereas *integrated* refers to studies that explicitly include EC variables as inputs in the assessment of ES. The review framework systematically categorizes the data from the studies using established taxonomies, ensuring the subsequent analysis directly addresses the overarching research question and the five specific sub-questions regarding EC integration, flow type, spatio-temporal dynamics, method classification, and condition variables.

The systematic review was conducted following the PRISMA 2020 methodology (Page et al., 2021), comprising the four core stages of identification, screening, eligibility, and data analysis. Given the rapid development and consolidation of the ecosystem services framework, we restricted the search to the last two decades (2005–2024) to reflect contemporary literature and assessment approaches. Searches were run on both Web of Science and Scopus databases to ensure maximum coverage of the scientific literature. Results were limited to English-language peer-reviewed journal articles, book chapters, and reviews. The search string, included in Supplementary Material 1, was constructed around four conceptual blocks: Urban, Ecosystem Services, Ecosystem Condition, and Method. (Fig. 1). It helped ensuring the retrieved studies develop or apply methods to evaluate urban ES while explicitly considering condition or biodiversity. Records from both databases were merged and de duplicated, resulting in an initial identification of 2836 papers.

**Figure 1.**
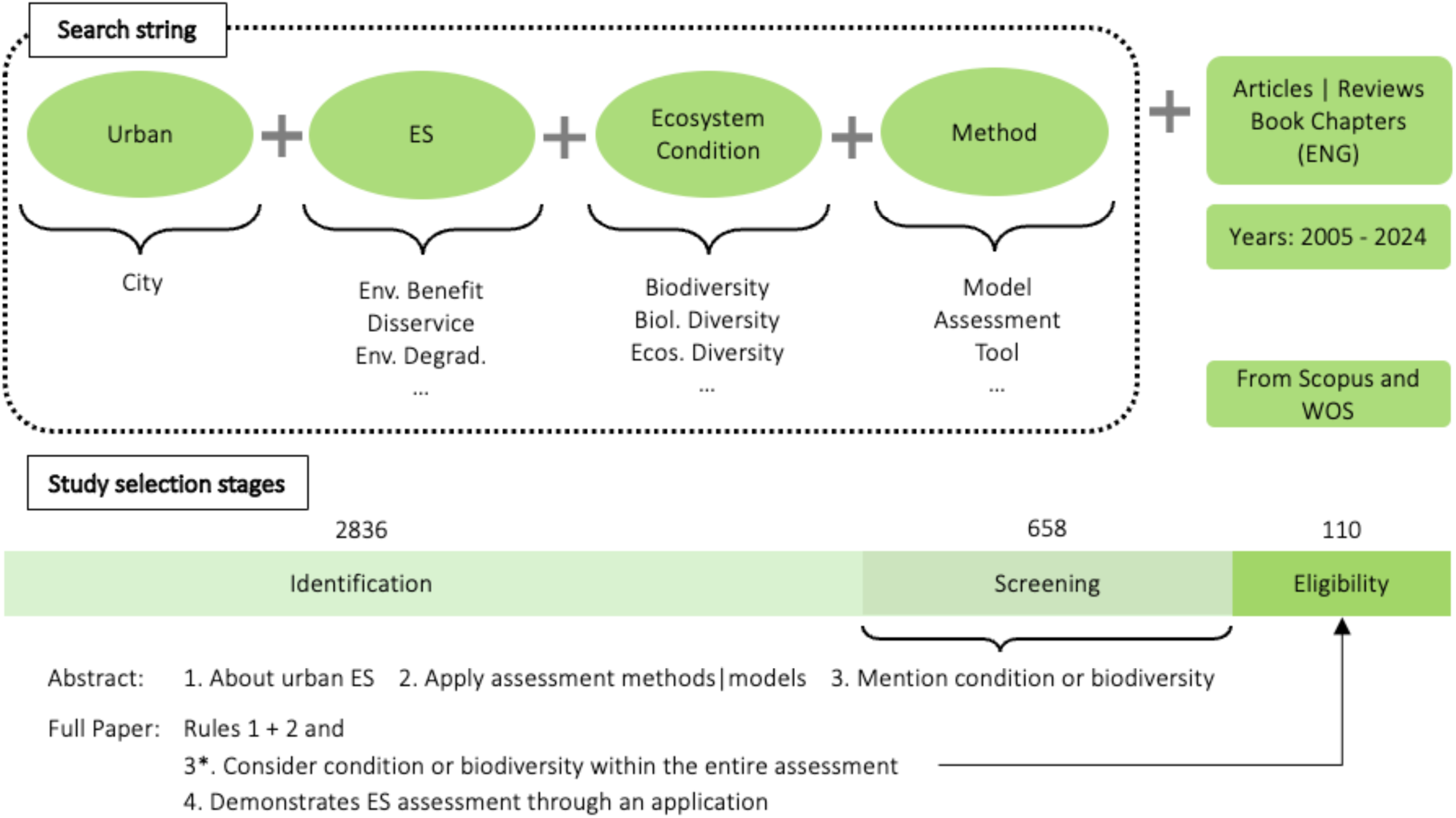
Definition of the search string and selection study stages of the literature review.

The study selection process followed the first three steps of PRISMA, as outlined in Figure 1. The initial screening was conducted at the title and abstract level. At this stage, papers were retained only if: (i) they included an ES assessment within an urban setting; and ii) mentioned ecosystem condition or biodiversity. When the inclusion of EC or biodiversity was unclear from the abstract, the paper was provisionally retained for full-text appraisal to minimize potential exclusions. This conservative approach resulted in 658 papers advancing to the full-text eligibility stage. During this stage, full texts were thoroughly examined. Eligibility criteria required that studies not only mentioned EC or biodiversity but also: (i) integrate EC variables into ES estimation (e.g., as predictors or state variables); or ii) assess ES and EC independently while explicitly coupling their analyses. In addition, only studies that demonstrated ES assessments through applications, such as case studies, scenario analyses, or virtual applications, were retained. This selection process resulted in 110 papers deemed suitable for the data analysis stage.

All retained studies were coded in a structured spreadsheet to extract and systematize the data relevant to our research questions. The coding framework was designed to capture three main dimensions: (i) the contextual features of each study; (ii) how ES were modelled and quantified; (iii) how biodiversity and EC were represented and considered into ES assessments,

We characterized the contextual features of each study by recording the following attributes: the spatial scale of application (site, neighborhood, city, region) of the ES assessment; the target urban ecosystem asset or nature-based solution assessed (e.g., street trees, parks, green roofs, urban wetlands); the presence of a real case study; and the country in which the ES assessment was conducted. This information helped us to understand the geographical coverage, the main ecosystem assets of interest and the typical scales of application represented across the reviewed studies.

To understand how ES were modelled and quantified, first we classified methodological approaches as biophysical, economic, or socio-cultural, and then coded the specific ES methods (e.g., process based models, macro-ecological models, hedonic pricing, participatory GIS) making use of the ESMERALDA typology (Brander et al., 2018; Santos-Martín et al., 2018; Vihervaara et al., 2018). ES classes were classified according to the SEEA-EA reference list to maintain consistency with accounting-ready service categories. In addition, we recorded the following features, considered relevant to address our research questions:

i. whether ES methods in a paper were applied in an integrated form (i.e., several methods concatenated to estimate values for one or more ES classes) or in a stand-alone form (i.e., a single method or multiple methods applied independently);
ii. the input and output variables used;
iii. the type of outputs obtained (quantitative/semi-quantitative/qualitative);
iv. whether ES methods were spatially explicit;
v. whether ES methods accounted for temporal dynamics in ES supply;
vi. whether models estimated ES potential or actual flows (i.e., ES demand/use was represented implicitly or explicitly); and
vii. whether sustainability thresholds were specified or discussed in any way.

This coding of ES classes and methods helped us to understand if ES assessments are done as a unicum or usually through standalone methods, and whether studies estimate potential vs. actual flows and consider sustainability of ES flows (RQ2), and if methods are spatially explicit and/or dynamic (RQ3).

To understand how biodiversity and EC were represented and considered, we coded whether studies explicitly integrated these variables into ES models or considered them only in coupled analyses (RQ1). For each study, we recorded the specific EC variables used and classified them in two ways. First, we applied the SEEA-EA EC Typology, which categorizes EC variables into abiotic (physical, chemical), biotic (compositional, structural, functional), and landscape characteristics. Second, we classified EC variables according to the Essential Biodiversity Variables (EBVs) framework, noting that this classification excludes abiotic variables and some biotic and landscape variables. This dual classification enabled us to interpret the use of condition variables under both international standards. Furthermore, we recorded whether variables referred to characteristics at the genetic, species, or ecosystem level of biodiversity, and whether these variables represented local, regional, or global values, that is, whether data were collected at the study scale or derived from regional or global averages. This coding of biodiversity and EC variables helped us to identify, across ES classes and method types the consideration of EC variables (RQ4), as well as the spatial accuracy of the EC data, and to clarify which dimensions of ecosystem condition are most commonly represented in urban ES assessment methods and models (RQ5).

All fields and coding rules are documented in a structured spreadsheet provided as Supplementary Material 2, and the related README file (as Supplementary Material 3) detailing the codebook, decision rules, and variable definitions to ensure transparency and reproducibility.

## 3. Results

The review encompassed 110 papers, representing only 3.9% of the 2836 initially screened. 108 of these papers contained case studies, while the remaining two applied ES assessment methods in virtual cases (Giedych and Maksymiuk, 2017; Pretzsch et al., 2023). Geographically, the studies span over 36 countries worldwide, with nearly half (17 countries) featuring only a single case study (Figure 2). A large number of studies are from Western Europe and the Far East; in particular, China, United Kingdom, Germany, and Italy, each contributed more than 10 case studies. In contrast, vast regions such as Africa and the Middle East are notably underrepresented. Regarding the urban ecosystem assets or nature-based solutions assessed, most studies could not be assigned to a single specific asset or solution type. Many articles focused on urban trees (29%) and urban green spaces (22%). However, studies in the latter category typically aggregated multiple asset types—such as various kinds of water features, trees, shrubs, and herbaceous plants—without distinguishing their individual contributions.

**Figure 2.**
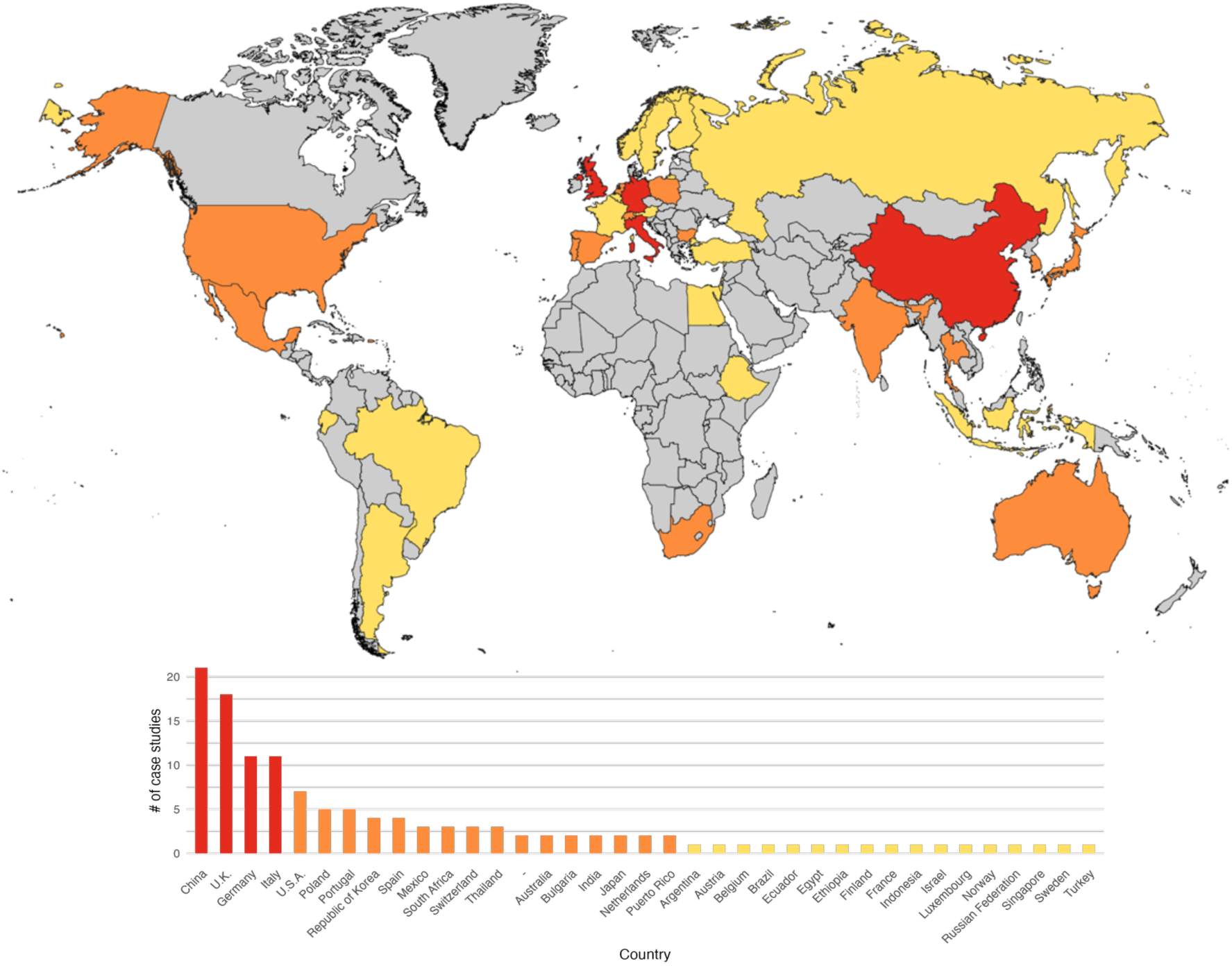
Geographic distribution of case studies that consider ecosystem condition within urban ecosystem service assessments. The world map displays the countries where case studies were conducted, and the accompanying bar chart shows their frequency. A three-color scale is used in both visuals: yellow indicates countries with one case study, orange represents those with two to nine, and red denotes countries with ten or more.

### 3.1. The “How”: Operationalizing ecological condition in urban ecosystem services assessments

Methods were applied integrated in just 30 articles; the rest of the studies (80) were applying single method or multiple methods but standalone (Fig. 3a). Regarding the consideration of EC in urban ES assessments (RQ1), the vast majority of the studies (96 papers out of 110) integrated EC as inputs or state variables within ES assessment, indicating that condition functioned as a core component of assessment methods. Conversely, only 14 papers merely coupled EC and ES assessments, and 23 did both (Fig. 3a). When EC integration occurred, variables were most frequently from the ecosystem level (86 papers), followed by the species level (66 papers), with 42 studies incorporating both levels. The genetic level was not addressed in any of the studies (Fig. 3a).

**Figure 3.**
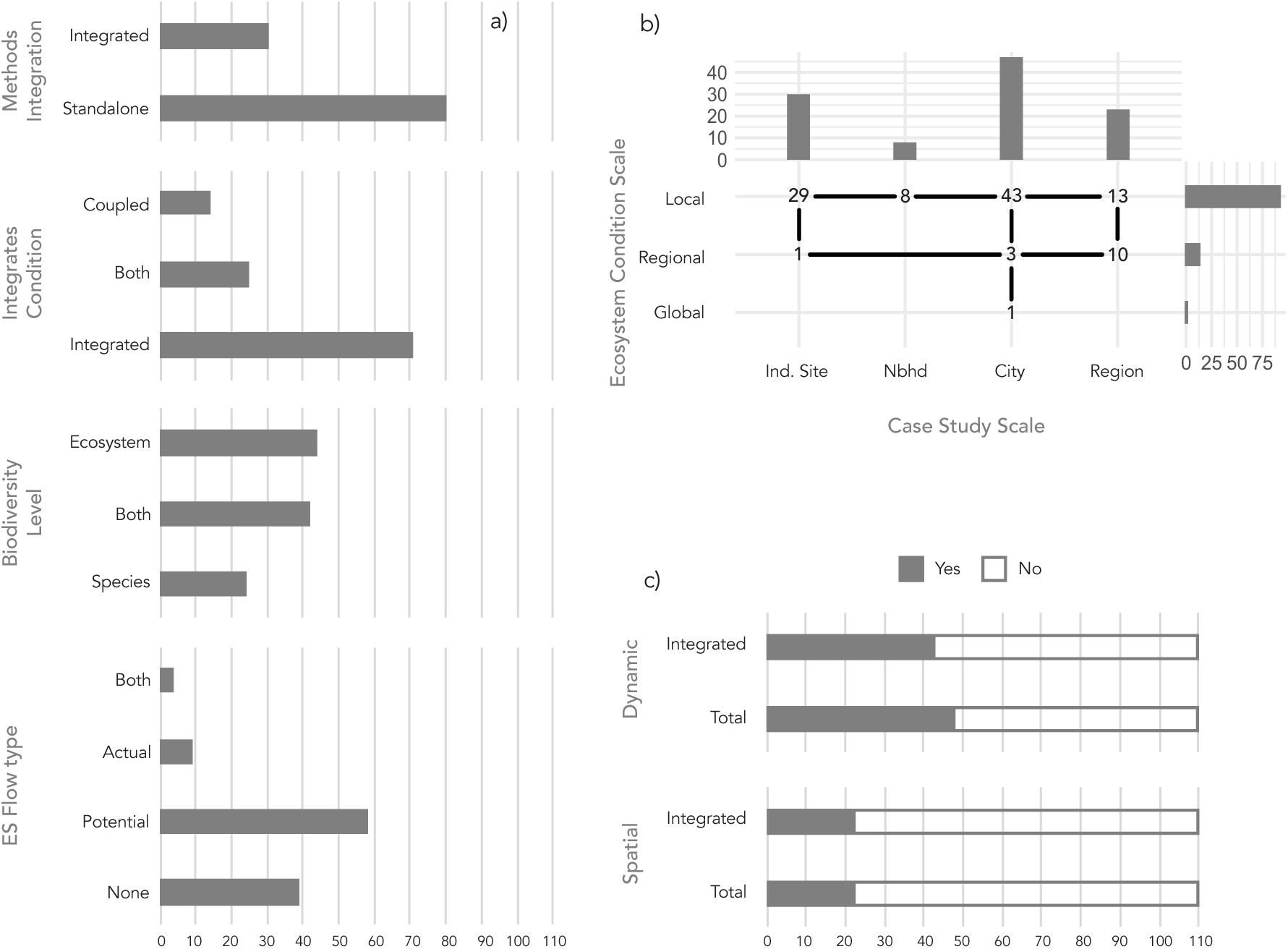
Overview of the bibliometric analysis of the full dataset. a) Number of articles in each of the four classification categories: Methods Integration, Ecosystem Condition Integration, Biodiversity Level and ES Flow type. b) Correlation between the biodiversity scale of the variables used (local, regional, or global) and the scale of the case study. Numbers within the matrix indicate how many articles fall within each combination, while the bar plot indicates the total number of papers per single category. c) Proportion of articles that are dynamical and spatialized, further filtered by Ecosystem Condition Integration.

In terms of whether ES assessments that consider EC estimated potential or actual flows, or informed about sustainability of the flows (RQ2), the analysis revealed that 39 studies did not consider a flow at all. This means that ES outputs were in qualitative or semi-quantitative form not assimilable to an ES flow. Among the studies estimating ES flows the majority (58 papers) focused on potential flows, while only 9 papers estimated actual flows, and just four papers considered both potential and actual flows within the same study (Fig. 3a). Among these four studies, three (Belaire et al., 2022; Dallimer et al., 2015; Dennis and James, 2017) studied potential flows for regulating and provisioning ES using biophysical models, often integrating EC variables like species richness or biomass density, while actual flows were consistently studied for cultural ES and were based on socio-cultural methods that did not integrate EC. Yan et al. (2024) in contrast assessed an actual flow for carbon sequestration in urban wetlands, incorporating real biophysical process data, such as photosynthetically active radiation and NDVI, thereby focusing on the actual production and realized service provision rather than potential capacity.

Two studies considered sustainability thresholds (Dallimer et al., 2015; Li et al., 2023), and other three discussed about sustainability implications (Elliot et al., 2022a; Perschke et al., 2023; Pretzsch et al., 2023). Four of these studies examined the balance between local ES supply and demand (Elliot et al., 2022a; Fanfani et al., 2022; Li et al., 2023; Perschke et al., 2023). In Li et al. (2023), sustainability was assessed by comparing ES potential with ES demand to classify systems as balanced, surplus, or deficit, with changes in ES potential linked not only to land use and cover but also to EC variables. Similarly, Fanfani et al. (2022) defined thresholds based on local food supply sufficiency, contrasting production capacity with per capita consumption. Elliot et al. (2022a) analyzed ES flow deficits within urban systems by comparing local supply to demand covered by both local and remote ecosystem assets, incorporating EC variables locally and estimating remote effects from land use/cover changes. Their study stands out as one of the few urban ES assessments to integrate EC while assessing transboundary ES flows and global sustainability implications. Pretzsch et al. (2023) approached sustainability differently, focusing on maintaining long-term ES supply through balanced urban tree stocks. Their work showcases how tree age structure, growth, and socio-ecological dynamics (e.g., natural mortality, harvesting, and safety-related removals) shape ES supply of urban forests over time, and how EC attributes such as leaf area, crown size, and tree size influence supply in non-linear ways. In synthesis, review outputs show that a few urban ES assessments incorporating EC, use it to understand whether ES flows are sustainable or not, when they do, most assess it indirectly by examining the balance between supply and demand or how condition characteristics and socio ecological dynamics of ecosystem assets underpin sustainability of long-term ES flows.

Looking at Figure 3b, we can see the relation between the scale of the case study and the scale of the EC variables incorporated. Theoretically, if the studies focused in an Individual Site, the EC should be local, while on the contrary if the article focuses on the urban region, it should estimate EC variables values at the regional level. Looking at the table, while it is mostly true that for most fine-scaled studies the variable used are usually local, when we move to the urban region 10 out of 23 times the study is informed with EC at local scale. Focusing on the EC scale, most of the variables represented ecosystem characteristics at the local scale (85 papers) with only a few studies drawing on data derived from regional (14) or global (3) scales. The inclusion of regional-scale EC data was consistent with studies assessing ES across multiple urban contexts or large metropolitan regions, such as in several cases from China (Liu et al., 2023; Tang et al., 2020). The combination of EC variables from different spatial scales was rare; only one study integrated local and global variables (Muluneh and Worku, 2022). This pattern confirms a generally good integration of EC when the assessment scale is fine (local), but the persistent reliance on local-scale EC variables in almost half of the assessments targeting the urban regional scale may indicate a spatial mismatch when scaling up to larger, more complex contexts.

Regarding the spatial and temporal explicitness of urban ES assessments (RQ3), as visualized in Fig. 3c, approximately 45% of all studies included spatially explicit methods. This proportion remains similar (43 out of 98 studies) when considering only those integrating EC into the urban ES assessment. In contrast, the incorporation of temporal dynamics is substantially lower, with only about 20% of the papers (22 out of 110) applying dynamic methods. A share that is maintained when focusing only on studies integrating EC into ES assessments (Fig.3). 9 out of 22 articles that appear to be dynamic are applying i-Tree model. Among the rest, Melliger et al. (2017) studied the soil quality regulation provided by tree litter and microbial activity, focusing on monthly differences influenced by the changing climate, while Pretzsch et al. (2023) emphasized the importance of dynamic methods, as EC– ES relationships are often non-linear. Just 9 studies (Elliot et al., 2022a; Krivtsov et al., 2022; Manes et al., 2016, 2012; Prigioniero et al., 2022; Silli et al., 2015; Sudarma et al., 2024; Yan et al., 2024; Yang et al., 2018), integrating EC into ES assessments, combined both spatially explicit and temporally dynamic methods. Among these, Manes et al. (2016) studied air filtration, specifically PM_10_ and O_3_ removal, accounting for process-based changes of the trees, in particular of Leaf Area Index, while maintaining the spatial explicitness of the distribution of the trees in the environment. Sudarma et al. (2024), accounted for carbon sequestration studying NDVI of the tree from satellite images, thus for large areas with changes from year to year. These results suggest that, although many urban ES assessments integrating EC as inputs in methods adequately address spatial dimensions, a substantial proportion still fail to capture critical temporal changes in EC and service flows, and even less capture spatiotemporal dynamics.

### 3.2. The “To What Extent”: Prevalence ecosystem condition across methods and services

Regarding the integration of EC by ES group, class and type of assessment method (RQ4), the overall distribution of studies across the three ES groups shows that regulating ES are the most frequently assessed (91 studies), followed by cultural (37) and provisioning services (29). As expected, biophysical is the most prevalent methodological approach (101 studies), far surpassing socio-cultural (15) and economic (15) approaches.

A closer examination of studies integrating EC variables by ES class reveals a highly uneven distribution (Table 1). Carbon sequestration and storage is the most commonly assessed ES class (51 articles), followed by air filtration (33), local climate regulation (29), and habitat maintenance (27). Within cultural ES, recreation-related services (25) is the most common ES class, followed by visual amenity (16), education and research promotion (7) and spiritual, artistic and symbolic value (6). For provisioning ES, crop provision (17), and water supply (12) are the most studied, while other classes appear in only one or two studies. Overall, many ES classes across all groups are represented in very few studies; despite identifying 29 ES classes in total, only 12 are considered in 10 or more studies.

**Table 1.**
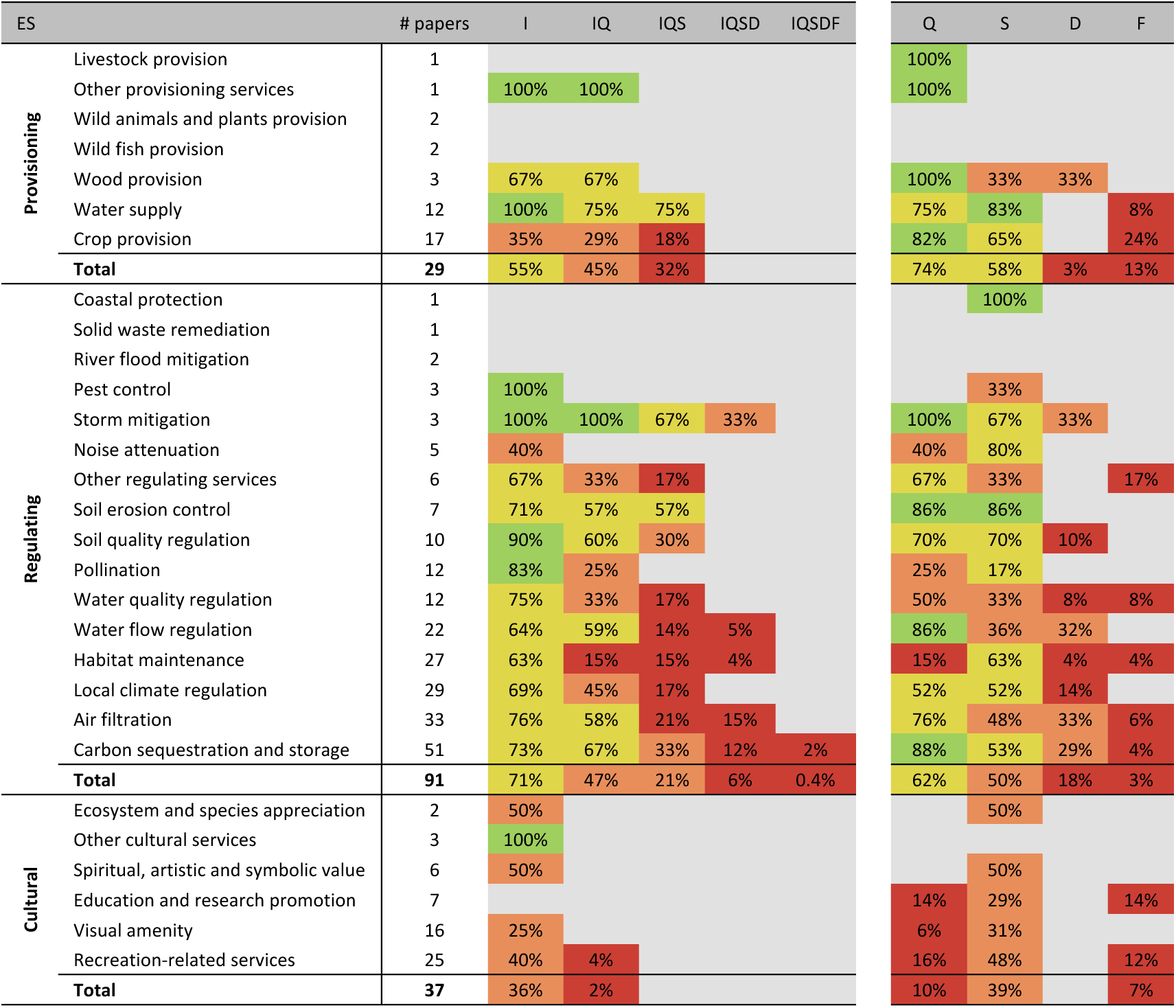
Share of papers for each ES, filtered for methodological characteristics. The specific characteristics used as filters are: I for being integrated, Q for quantitative results, S for spatialized, D for dynamical and F for Actual flows. Potential flow has not been considered as it does not modify the number of articles retained after being filtered for spatial explicit and dynamical. The colors are moving from green to red, as the proportion of articles in which that ES continues to be included (Green being above 80%, yellow between 80% and 50%, orange between 50% and 20% and red lower than 20%)

When disaggregating each ES class by specific methodological characteristics, namely whether methods are integrated (as opposed to standalone), whether outputs are quantitative, whether they exhibit spatial explicitness or temporal dynamism, and whether outputs are assimilable to actual flows, the unevenness of the distribution becomes even more pronounced (Table 1). Filtering only by integrated methods and quantitative outputs results in a minor loss of studies; however, when restricting the selection to studies that are simultaneously integrated, quantitative, spatially explicit, and dynamic, only five ES classes remain—all of them regulating services. Finally, when filtering for studies that account for actual flows it remains only one study accounting for carbon sequestration (Yan et al., 2024). For instance, while 37 papers use integrated methods for carbon sequestration, only 17 of these are quantitative and spatially explicit, and from these only 6 consider temporal dynamism. This unevenness in distribution is similarly evident when each methodological characteristic is considered individually. Temporal dynamism and outputs assimilable to actual flows are particularly underrepresented, with studies addressing only 9 ES classes in these cases. When applying the filters individually, most of the remaining ES classes correspond to regulating services, with a few exceptions such as recreation-related services, crop provision, and water supply.

In summary, the color-coded visualization in Table 1 highlights that while EC variables are already being integrated as inputs in urban ES assessments, this occurs predominantly for regulating services and only a few provisioning and cultural services. Moreover, when methodological features desirable in advanced or ecosystem accounting–oriented approaches are considered (e.g., spatial explicitness, temporal dynamism), the number of studies, and methods, decreases drastically, revealing that most studies remain non-dynamic and rarely assess actual flows.

When repeating the same analysis for assessment methods, we continue obtaining an unevenness in the results (Table 2). Within biophysical approach, process-based (43 studies), statistical (35), and spatial proxy methods (32) are the most frequently employed. Each single method for economic and socio-cultural approaches appears in less than 10 studies, with a maximum of 8 applications for market price and 5 for preference assessment. This is an expected result, since it makes sense that EC are integrated as inputs of biophysical methods and, in many cases, economic and socio-cultural methods are used in later stages of ES assessments, not including directly EC variables as inputs but ES as biophysical outputs.

**Table 2.**
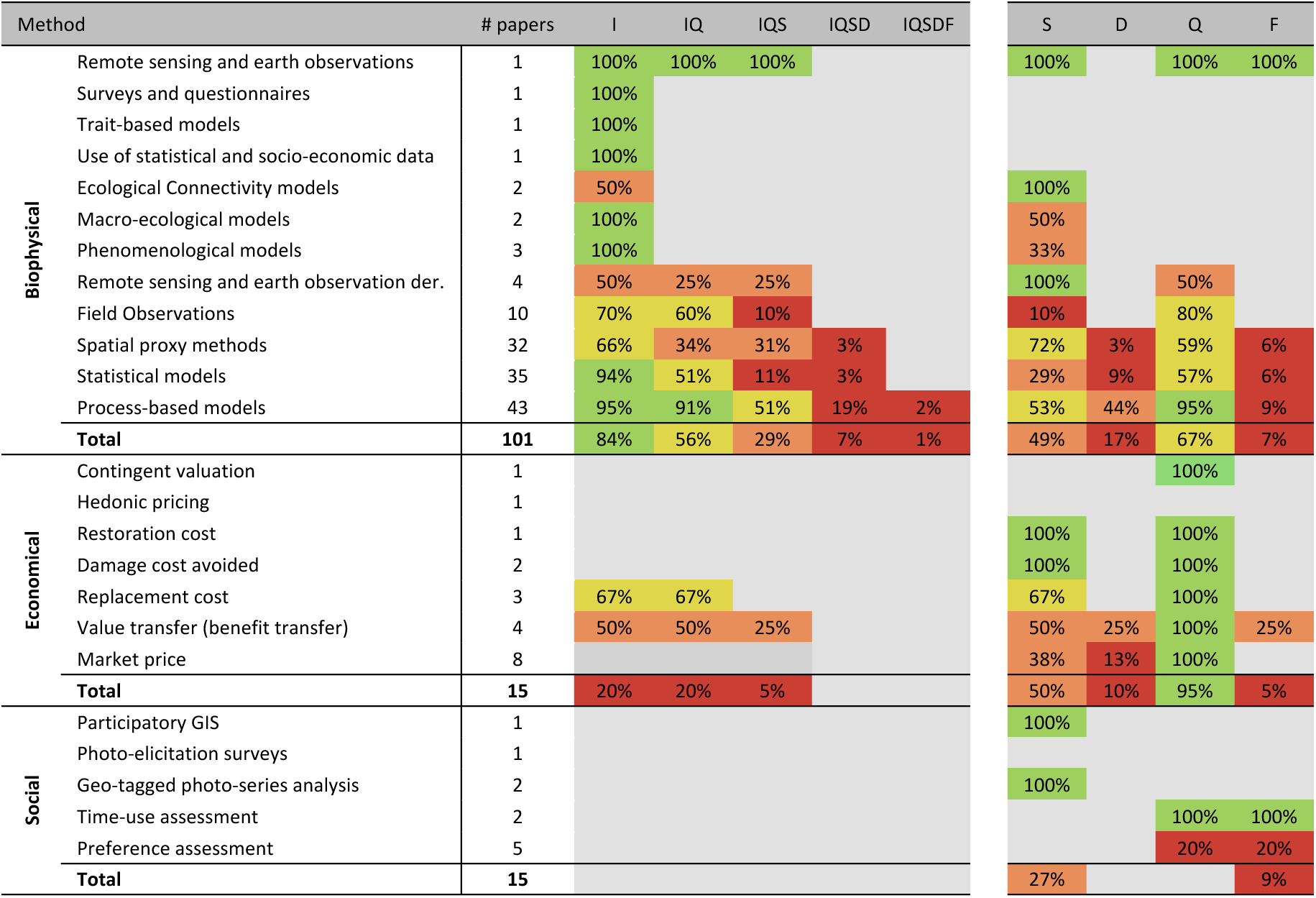
Share of papers for each Method, filtered for methodological characteristics. The specific characteristics used as filters are: I for being integrated, Q for quantitative results, S for spatialized, D for dynamical and F for Actual flows. Potential flow has not been considered as it does not modify the number of articles retained after being filtered for spatial explicit and dynamical. The colors are moving from green to red, as the proportion of articles in which that Method continues to be used (Green being above 80%, yellow between 80% and 50%, orange between 50% and 20% and red lower than 20%)

As per ES classes, when filtered by the same characteristics previously used, the number of methods applied decrease drastically, with only three types of methods that remain after filtration for integrated, quantitative, spatially explicit and dynamical (the same nine articles that are both spatial and dynamical); after filtering for actual flow it remains just the paper of Yan et al. (2024). Similarly, when considering methodological characteristics individually, the unevenness is only particularly relevant when filtering by temporal dynamism and actual flows, and not for all types of method. For example, spatial proxy methods and statistical models considering temporal dynamism are only used in one (Yang et al., 2018) and three studies respectively (Melliger et al., 2017; Sudarma et al., 2024; Torngern and Leksungnoen, 2020), but more than 50% of the studies using process-based models are still retained.

In summary, similar to the case of ES classes, the color-coded visualization in Table 2 highlights that while EC variables are already being integrated as inputs in multiple ES assessment methods, when methodological features desirable in advanced or ecosystem accounting–oriented approaches are considered, the number of studies and methods decreases drastically, but in this case this is also a consequence of very few methods (4) being applied in more than 10 studies.

### 3.3. The “To What Extent”: Nexus between methods, services, condition categories, and metrics

An analysis of interconnections between ES classes, methods, and types of EC variables (Fig. 4) for both the SEEA-EA and EBV typologies (Fig. 4a,b) identifies which types of EC variables are typically included in urban ES assessments (RQ5) and reveals how they relate to specific ES classes and methodological approaches.

**Figure 4.**
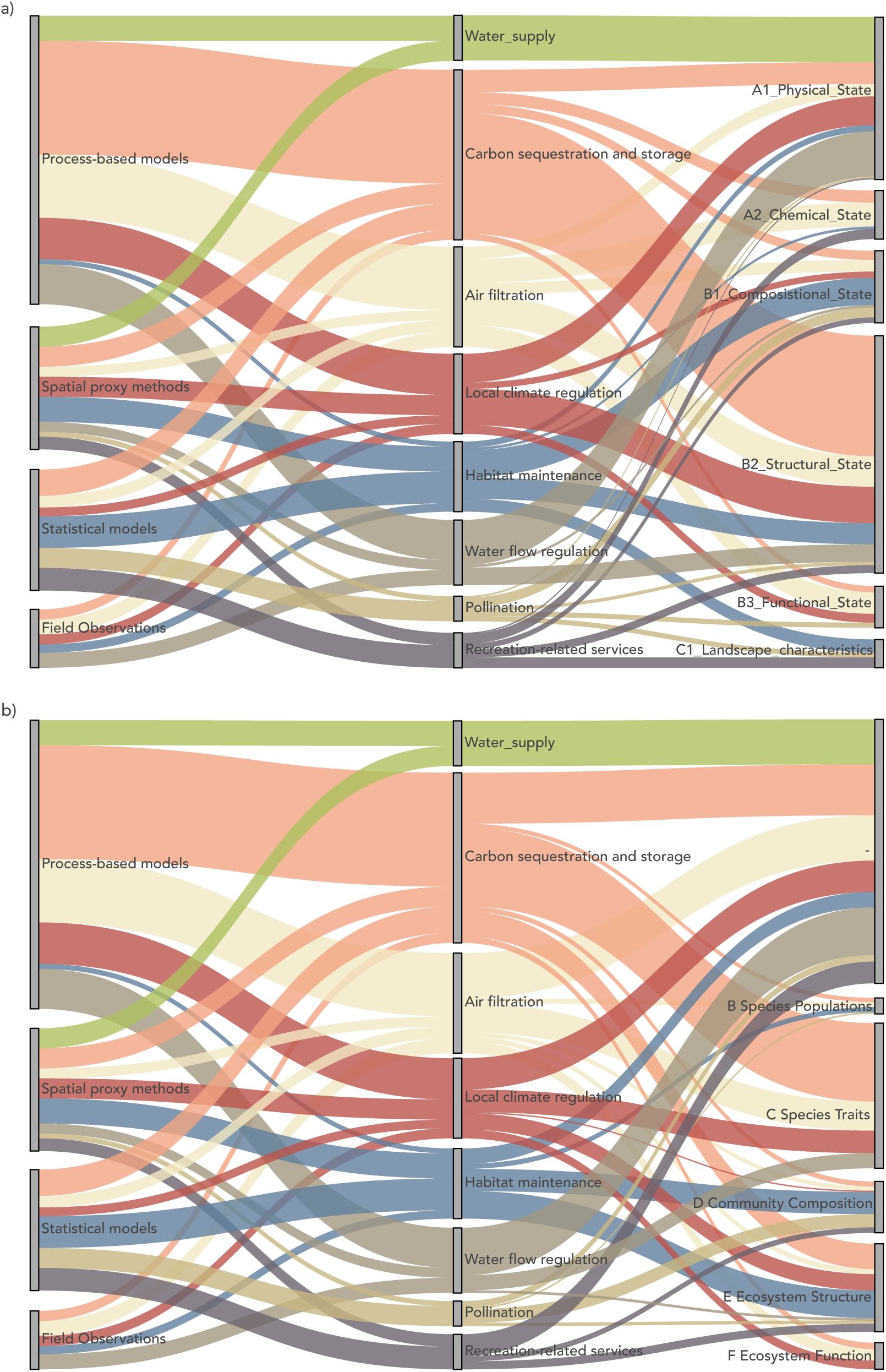
Sankey diagram of article connections between Methods, ES and a) SEEA EC or b) GEOBON EBV. Both Methods and ES are retained when are presents in more than 10 articles which limits the selection to 6 methods and 8 ES. Each color represents a different ES, to better visualize the connections. The connections are weighted according to every ES present in a paper.

While many ES classes exhibit complex patterns across methods and condition variables, several distinct trends emerge from Figure 4. For instance, carbon sequestration and storage is primarily examined using process-based models. According to the SEEA-EA EC typology, this service is mainly estimated using EC variables from structural state characteristics, and species traits according to the EBV framework, reflecting dependence on vegetation structure and biomass inputs. As another example, water supply is exclusively linked to abiotic factors, specifically physical state characteristics (SEEA-EA), and is never assessed using variables explicitly connected to biodiversity, such as those represented in EBV categories. The same, but less strict, pattern is observed for water flow regulation, where ES assessments rarely employ biotic factors, particularly those represented in EBV categories, despite the expectation that they would also play a relevant role in the supply of this service. In contrast, air filtration is frequently assessed using process-based models (more than half of the studies) and shows a strong dependence on EC variables from structural and functional state characteristics, which may indicate an emphasis on functional traits relevant to pollutant capture. Among all ES classes, habitat maintenance stands out by the fact that EC variables belonging to landscape characteristics (SEEA-EA) and ecosystem structure (EBV) are mostly used in its assessment, underscoring the key roles of spatial composition and configuration of ecosystem assets for the supply of this service. Finally, recreation-related services are also distinctive, as their assessments predominantly rely on statistical models, reflecting the use of correlative approaches that link the presence of green space with demographic data.

The relationship between both EC classifications, SEEA-EA and EBV, is clarified in Figure 5a, further informing about the types of condition variables included in urban ES assessments (RQ5). The diagram illustrates that structural state characteristics followed by physical state characteristics are largely the most frequently used EC category from SEEA-EA typology. Among the specific EBV categories, morphology and habitat structure are among the most commonly used. However, a notable proportion of variables that can be classified with SEEA-EA typology are not linked to specific EBV. This reflects the importance of abiotic factors as EC variables and suggests that a portion of biotic factors of relevance may not be captured within the biodiversity categories of EBV. In addition, mostly functional state characteristics show connections to primary and secondary productivity (EBV), indicating that when function is the focus, the proxies relate mostly to the overall photosynthetic output and growth of the ecosystem.

**Figure 5.**
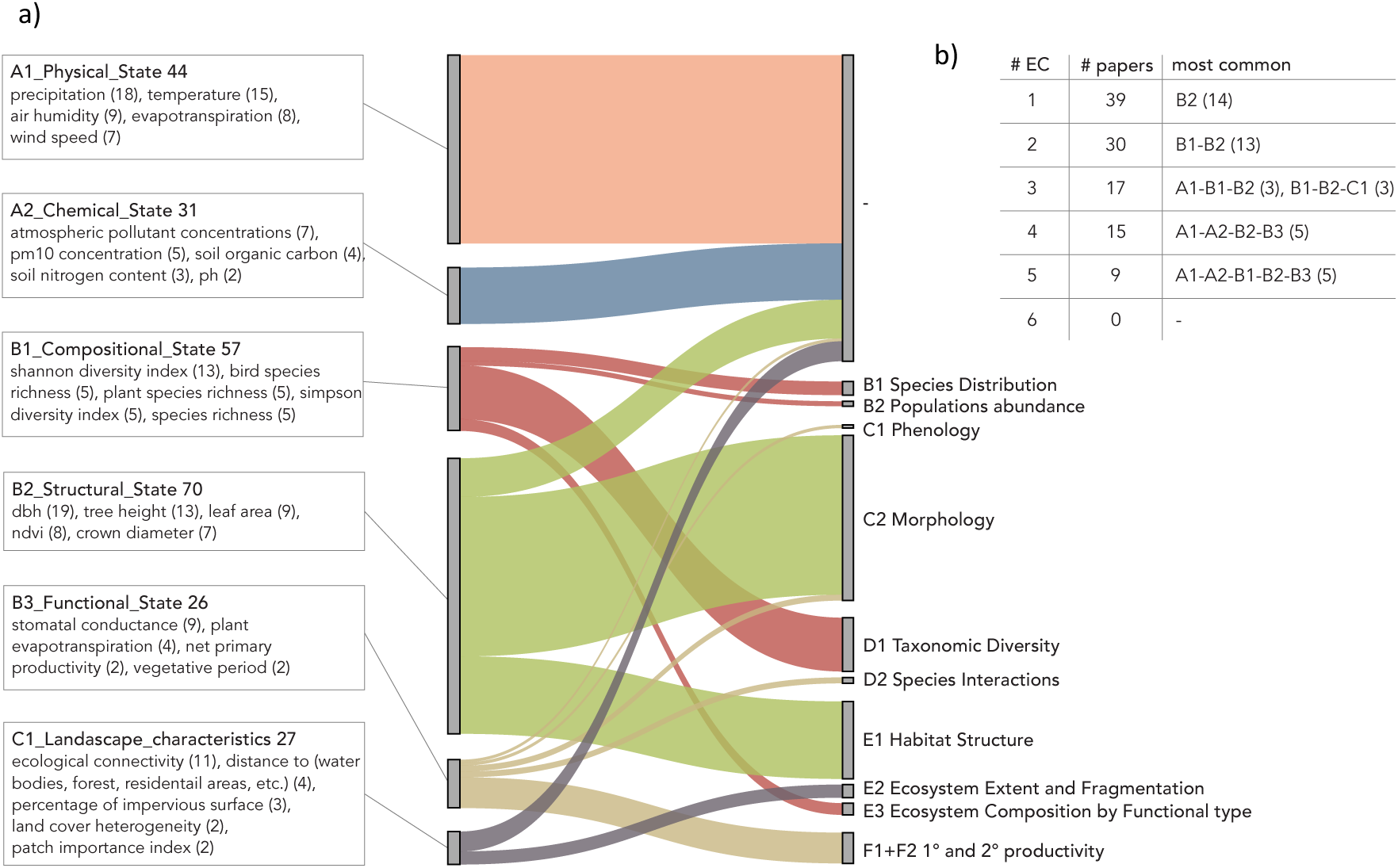
a) Sankey diagram representing the connections between EC and EBV. The colors represent the EC, and the weighting of the connections is proportional to each ES present in a paper, as in Figure 4. For each EC the most common specific variables used for each SEEA-EA EC categories; the number after the EC refers to the number of paper in which the specific EC is used (not filtered for integration); the number in bracket after the specific variable refers to the number of paper in which that specific variable is used. b) Count of the papers in which 1 or more EC are used at the same time, with the most used combination of EC for each number of EC present.

To ground the relational patterns between both EC classifications to specific variables, Figure 5a also details the recurrent variables used as proxies within each category. This breakdown reveals how variables often bridge or fall outside the boundaries of one of the classifications. For instance, within the SEEA-EA Physical State (A1) category, precipitation and temperature are the most frequent variables. Their high usage is likely due to their wide accessibility and ease of measurement across various settings; but being abiotic they do not fall under any specific EBV category. Figure 5a also shows that frequent EC variables are not always directly integrated in ES calculations, but just coupled. This is the case of the Shannon Diversity Index, which is the most common variable for SEEA-EA Compositional State (B1) category and EBV Taxonomic Diversity (D1) category. Notably, despite being incorporated in 13 studies, only three of them integrated the index directly into the ES calculation as a proxy for habitat maintenance, pollination, and recreational-related services (Hernández-Rivera et al., 2024; Liang et al., 2023; Prigioniero et al., 2022). This gap may be explained because the relationship of this EC variable with ES supply is not always simply and direct. As an opposed example, under the SEEA-EA Structural State (B2) category and EBV Morphology (C2) category, diameter at breast height and tree height are variables largely utilized in urban ES assessments, and directly included in ES calculations, mainly to quantify carbon sequestration and storage. As a middle case situation, just one recurrent EC variable of SEEA-EA Landscape characteristics (C1) category, proxies of ecological connectivity, is also included in an EBV category (Ecosystem extent and fragmentation – E2), and in some studies integrated in ES methods for the calculation of habitat maintenance and water supply (Oertli et al., 2023; O’Keeffe et al., 2022; Ranta et al., 2021; Raquel Calapez et al., 2023). Instead, other common variables representing attributes of the landscape mosaic are not directly captured within the EBV classification.

Figure 5b addresses the extent to which urban ES assessments consider EC comprehensively, shown by the number and specific combination of EC categories used simultaneously. The results show that most studies use only one or two categories (35% and 27%, respectively). The use of three or more categories is less common, with the proportion dropping to 15% for three and 14% for four. Notably, comprehensive integration is rare; only 8% of the studies (9 of 110) used five categories, and no study incorporated all six. Comprehensive studies often targeted multiple ES or large areas; for instance, Lessi et al. (2024) studied a variety of ES with semi-quantitative output in São Carlos Municipality, and Yang et al. (2018) employed an emergy assessment (Odum, 1996) using various EC as spatial proxies.

In synthesis, the findings presented in Figures 4 and 5 confirm that overall, and across ES classes and assessment methods, structural, physical, and easily measurable EC variables are strongly preferred to be directly integrated into urban ES methods and models.

## 4. Discussion

### 4.1. Interpreting the “How”: progress in methodological operationalization vs. shortfalls in capturing complexity

Our findings show clearly that the majority of the studies retained opt for integrating EC variables as functional components of urban ES assessments and not just merely coupling the EC and ES assessments. This points out to EC as a crucial biophysical variable in ES methods, moving away from simple post-hoc correlations. However, this progress in terms of methodological integration of EC is not mirrored in the capacity of these urban ES assessments to capture comprehensively the socioecological complexity, particularly concerning the estimation of ES supply flows and their sustainability and their temporal dynamism.

Around one-third of the studies estimated ES using qualitative or semi-qualitative outputs. As a consequence, these methods cannot be used in urban ecosystem accounts, because their outputs cannot be assimilable to supply flows. At first, this finding may not seem a real limitation, since not all urban ES assessments should be useful to inform methods valid for ecosystem accounting. However, it does indicate that many methods still rely on ad-hoc (semi)qualitative scales that may hinder comparability across studies and restrict integration into broader quantitative analyses, such as cost– benefit assessments.

Among the studies where ES were assimilable to flows, our findings show that very few estimated actual flows. In fact, in those studies there was a clear disconnect between EC and actual flows, since EC integration was systematically used to model the potential supply of regulating and provisioning services, while actual flow was estimated for cultural services using socio-cultural metrics that excluded EC variables (Belaire et al., 2022; Dallimer et al., 2015; Dennis and James, 2017). Even fewer studies used EC to reflect on the sustainability of ES flows, and they did it examining the balance between supply and demand (Li et al., 2023) and comparing the provision of ES in different years (Dallimer et al., 2015). This suggests an indirect assessment rather than the integration of critical sustainability thresholds within urban ES assessments. It may therefore highlight a major limitation: despite the coupling or integration of EC variables, they are not used to inform whether an ES mismatch exists that could lead to degradation of urban ecosystem assets in terms of their intrinsic, relational, or instrumental values. For example, only a few studies integrating EC variables provided information on whether the ecosystems’ capacity for the assessed ES was being exceeded, which is expected to lead to degradation of ecosystem quality and long-term ES supply. Consequently, findings indicate that the integration of EC variables into urban ES assessments is still not translating into an improved understanding of the sustainability of ES supply or of whether the quality of ecosystem assets is being effectively preserved.

The scarcity of temporally dynamic methods (only 20%) also emerged as a relevant limitation, as EC–ES relationships are frequently non-linear and time-dependent (Pretzsch et al., 2023). Most urban ES assessments integrating condition still treat abiotic, biotic and landscape characteristics as static values, limiting the utility of the assessments for adaptive management or forecasting the effects of climate change or long-term management interventions. In those cases, with strong seasonal variation in ES supply and demand, it may not permit understanding if urban ES are supplied when demand exist or it is high, therefore effectively contributing to generate a human well-being benefit.

As a positive outcome, our findings show a quite coherent relation between the spatial scale of the case study and the spatial scale at which data for EC variables are gathered. In fact, studies doing ES assessments at fine scales (individual site, neighbourhood) mostly use local data for EC variables (e.g., site-specific tree or soil measurements), with regional averages used just for assessments at city or urban region level.

### 4.2. Interpreting the “To What Extent”: a skewed landscape across services, methods, and metrics

Our findings confirm that the integration of EC variables into ES methods is heterogeneous across ES classes and types of methods, with cultural and provisioning services remaining behind. The findings also suggest that in some cases EC integration or coupling is not incentivized and operational feasibility is prioritized over a comprehensive consideration of different types of condition variables in urban ES assessments.

In terms of ES methods, the dominance of the biophysical methodological approach was expected because ES are usually first calculated in biophysical units. Then, the consideration of EC may not be needed when economic and social methods are successively applied. In fact, from the cascade framework, the biophysical estimation may be implicit and followed by an estimation in monetary or social values (e.g. quality of life scores) of the contribution of ES to human well-being. Similarly, the SEEA-EA framework explicitly calls for estimating ES physical accounts before monetary accounts. Thus, EC is expected to be considered mostly in biophysical methods.

Regulating ES, in particular carbon sequestration, air filtration, and local climate regulation, was the ES category where EC was more frequently integrated and which included more temporally dynamic methods. From one hand, this may be explained from the fact that regulating ecosystem services (ES) constitute the predominant category (91 studies) in our review. This is an expected outcome, as recent reviews on urban ES and NBS highlight regulating services as the most frequently assessed category in urban areas (Babí Almenar et al., 2021; Li et al., 2025). On the other hand, these ES classes depend directly on measurable biotic and abiotic characteristics (e.g., leaf area, biomass, structural complexity, air pollution concentration), already incorporated in well-known modelling suites, such as i-Tree, and widely applied process-based models. Some of these properties, such as the leaf area and air pollution concentration, are known to vary strongly at seasonal or even hourly time scales which also explain why temporally dynamic methods are more recurrent.

The lower representation of provisioning and cultural ES classes, besides reflecting that these are categories less studied in urban areas, may also be partly associated with three different causes. For provisioning ES, such as crop provision, wood provision or water supply, because they already have markets, services are not a positive externality, and there are well-developed inventories compiled regularly on yields or volumes produced. For instance, in our review Fanfani et al. (2022) relied on such inventories to estimate the local crop provisioning capacity, implicitly assuming that the local supply of this ES class was sustainable. As a result, it becomes relatively easy to associate these estimates, typically expressed as averages within a given area, with land-cover classes or types of ecosystem assets, that is, just with ecosystem extent data. Instead, for cultural ES, the relationship between EC and ES supply may be less evident or more indirect, making EC integration more methodologically challenging. For example, EC variables, typically biophysical in nature, are less straightforward to integrate into models assessing human perceptions or the spiritual and symbolic values of urban ecosystem assets. As a third cause, for both provisioning and cultural ES, it is also difficult to disentangle the contribution of human artifacts and actions (e.g., availability of footpaths, benches, water points, human labor) from the contribution of the ecosystem asset itself in the ES supply. The above causes may limit the coupling or integration of EC in urban ES assessments for provisioning and cultural services, since the additional data requirements and modelling complexity may not be justified.

Our findings also revealed a heterogenous adoption of different types of condition variables according to both SEEA-EA and EBV typologies, characterized by a clear dominance of structural and physical state variables. This pattern is likely not coincidental but reflects a pragmatic alignment with data accessibility and the existence of established modelling suites already incorporating such proxies. For instance, variables like diameter at breast height of trees are not only easily acquired but also exhibit a direct, often linear relationship with services like carbon sequestration, facilitating their integration. Conversely, compositional and functional variables, while conceptually central for biodiversity and ES relationships, are less frequently used. The underutilization of these variable types may be partly attributed to the higher measurement effort they require and, critically, to their more complex, frequently non-linear and time-lagged relationships with ES flows. Consequently, while the prevailing use of easily measurable condition variables enables the calculation of many ES flows, it may fail to comprehensively capture the functional health and biotic complexity of urban ecosystem assets. This limitation ultimately may restricts achieving a deeper understanding of how biodiversity, and more broadly ecosystem quality, influences ES supply in urban systems.

### 4.3. Practical implications for service modelling, urban greening actions and urban ecosystem accounting

The widespread integration of physical and structural state EC variables has enabled the development of accessible single-service models, even in the form of well-known suites like i-Tree or SWMM. While these models facilitate straightforward urban ES assessments for baseline conditions, their narrow focus limits their capacity to capture the full complexity of ecological processes that underpin ES supply. For example, while tree height (structural state variable) predicts carbon stock, the diversity of functional traits (compositional state variable) determines its resilience to pests or climate change (Díaz et al., 2013). This limitation is critical for two reasons. First, for certain services understudied in urban areas, like pest control, accurate modelling inherently requires compositional or functional EC variables, even for single-service models. Second, to predict the capacity of urban ecosystem assets to supply multiple ES under future scenarios, models must account for the interdependencies between services, which reinforces the need of integrating proxies from these less used EC classes. Incorporating such complexity is essential for improving the realism of long-term future ES estimations, particularly to account for risks from future recurrent stresses and shocks. Ultimately, evolving models in this direction is crucial for informing urban greening plans that aim to secure resilient ecosystem assets.

The current integration of EC variables into urban ES assessments is beginning to shape urban greening actions, primarily by reinforcing the need for fine-scale and continuous or regular monitoring of abiotic and structural biotic variables. In contexts like the EU, new regulations (e.g., the Nature Restoration Regulation) are accelerating this trend, prompting local and regional administrations to mainstream granular monitoring systems that offer a dual benefit: tracking asset quality and enabling more accurate ES estimation. However, this progress has critical limitations. First, it remains narrow; for example, comprehensive soil health monitoring in urban ecosystems still lags behind. This is a gap that the recent EU Soil Monitoring Law (EU Directive 2025/2360) aims to fill by requiring the monitoring of soil health beyond natural and agricultural soils and its evaluation based on physical, chemical, and biological condition proxies. Second, and more critically, the prevalence of static models in our review shows that available EC data have not yet been translated into mainstream dynamic ES forecasting. When combined with the rare estimation of actual ES flows, this means assessments partly fail to quantify the real-time contribution of greening actions to human well-being in baseline scenarios, not just in long-term future ones. Consequently, planning based on static potential ES supply risks overestimating benefits and misses opportunities to manage seasonal demand-supply mismatches. To effectively inform urban greening, the use of dynamic process-based models should be mainstreamed, allowing practitioners and decision makers to leverage fine-scale spatio-temporal EC data to simulate temporal changes in ES supply for more resilient outcomes.

The abovementioned progress and shortcomings have direct implications for advancing urban ecosystem accounts. The widespread use of abiotic and structural state variables, bolstered by new EU regulations (e.g., the EU amendment on environmental-economic accounts), will facilitate the implementation of initial urban condition accounts. However, our results indicate that accounts for compositional, functional, and landscape variables will likely lag behind. This expectation is reinforced by Nicholson Thomas et al. (2025), whose review found a severe scarcity of such condition variables in urban contexts, with not even a single structural or functional variable reported in multiple instances. Our findings also align with challenges identified by Babí Almenar et al. (2025) regarding the advancement of urban ecosystem accounts. We confirm, specifically within the context of urban ES assessments, the lack of reference condition thresholds and problems in assessing flow sustainability. Our review directly links these two issues, showing that the current integration of EC into urban ES models fails to incorporate condition thresholds to evaluate the sustainability of supply flows. Furthermore, we substantiate a shortcoming they highlighted for monetary asset accounting: the limited capacity of urban ES models to predict the supply of multiple, interdependent ES under future scenarios. Specifically, our findings highlight that assessments largely rely on standalone single service approaches that miss critical interconnections and the role of underlying ecological complexity. This inherently restricts the development of monetary asset accounts capable to reflect the cumulative value of multiple future ES flows beyond baseline scenarios.

### 4.4. Limitations and the novel synthesis of our review

Our systematic review provides a robust analysis of the consideration of EC into urban ES assessments, yet its specific focus introduces inherent limitations. Firstly, it provides a targeted view of the EC-ES nexus rather than a complete census of urban EC research. Broader reviews, such as those by Rendon et al. (2019) or Nicholson Thomas et al. (2025), which focus on condition variables across multiple ecosystems can be used to complement our findings. Secondly, our urban-specific scope may overlook relevant methodological advances from other ecosystems, such as in wetlands or agroecosystems (McLaughlin and Cohen, 2013; Rendon et al., 2022). However, the unique character of urban ecosystems (fine-grained, fragmented, a mix of intertwined natural and artificial surfaces, and of socio-ecological-technological nature) demands caution, as transferring EC-ES methods or findings often requires significant conceptual and operational refinement, particularly for ecosystem accounting purposes (Babí Almenar et al., 2025). A final, contextual limitation is that a substantial number of screened studies were excluded due to a complete lack of condition variables, underscoring that its integration is not yet standard practice. It is crucial to note, however, that our objective was not a bibliometric count of trends over the 20-year range of our review, but an analysis of methodological progress where EC coupling or integration in urban ES assessments *does* occur. This specific analytical focus leads to the core strengths of our work.

The primary strength of this review lies in its detailed methodological synthesis, which moves beyond acknowledging the consideration of EC to explicitly analysing its operationalization. We provide one of the first comprehensive linkages between specific ES classes (SEEA-EA classification), ES assessment methods (ESMERALDA classification), and distinct EC categories (SEEA-EA and EBV typologies) down to the level of recurrent individual variables. The use of established classifications for these three groups, rather than ad-hoc categories, enhances the generalizability of our findings. For instance, by employing both the SEEA-EA and EBV typologies, we offer a dual-perspective that highlights the importance of abiotic factors (often missed by biodiversity-centric frameworks like the EBV) and the strong preference for easily measurable structural variables.

Our results point that the conceptual dependency of ES on EC is widely accepted and the technical integration is advancing. Yet, the increase in literature volume does not seem to be translated into a deeper, more comprehensive understanding of the interrelation between ecosystem quality (condition) and ES supply flows. We identify specific gaps that may hinder advancing our understanding on EC-ES linkages: the stark underrepresentation of functional, compositional, and landscape state variables; the widespread neglect of the temporal dynamism of EC and therefore ES supply; the still rare assessment of actual ES flows; and the missed opportunity to leverage EC integration for understanding sustainability thresholds of ES supply. In summary, the added value of our work is in pinpointing that future progress depends on overcoming these specific gaps.

## 5. Conclusions

This review analysed how, and to what extent, ecosystem condition (EC) is considered in urban ES assessments. The findings indicate a clear operational integration of EC variables as functional inputs within ES models. However, this progress is markedly uneven. Integration is concentrated on regulating services, relies on static frameworks, focuses on potential over actual flows, and favors easily measurable structural and physical EC variables over functional, compositional, and landscape state variables. Consequently, while the conceptual EC-ES link is widely accepted, technically feasible, and practically progressing, it remains methodologically shallow, which limits the potential to increase our understanding of how biodiversity and ecological complexity underpin long-term ES supply.

These limitations constrain the practical value of EC integration into urban ES assessments. The lack of dynamic models and actual-flow assessments hinders the utility of urban ES assessments for the planning and adaptive management of urban greening, undermining the long-term value of these actions under future scenarios. Furthermore, they represent also a significant hurdle for advancing urban ecosystem accounting. Under the SEEA-EA framework, ecosystem accounting requires time series data on asset condition, defining reference condition thresholds and tracking actual flows over time. These elements are vital for assessing flow sustainability and diagnosing potential ecosystem degradation. Yet, a gap remains, as most urban ES assessments do not employ the kinds of models and methods needed to fulfil ecosystem accounting needs.

Despite these challenges, the review identifies a clear pathway forward. The established use of physical and structural state variables provides a practical foundation for immediate action. These established metrics can be standardized and leveraged to form the initial urban condition accounts and inputs in models suitable for urban ES accounts. Concurrently, future research must prioritize three frontiers: (1) developing dynamic models that link temporal changes in EC to ES supply; (2) leveraging EC integration to bridge the gap to actual service flows; and (3) incorporating a more holistic suite of functional, compositional and landscape state variables to better represent how ecological complexity influences ES supply. By building on current strengths to address these gaps, urban ES research can enhance the rigor and policy-relevance of its outputs, supporting the sustainable, evidence-based management of urban ecosystem assets.

## Supplementary Information

Supplementary Material 1. Complete search string used for the literature review

Supplementary Material 2. Data and metadata from the articles analysed in the literature review

Supplementary Material 3. README file with the instructions for the spreadsheet

## Acknowledgement

This work was supported by the National Biodiversity Future Centre (NBFC) project, funded by the European Union’s NextGenerationEU, National Recovery and Resilience Plan (NRRP), CN00000033, CUP, D43C22001250001.

