## Supplementary material for "Do urban ecosystem service assessments account for ecosystem condition and biodiversity?": SM1

### Search string

WOS search:

TS = (("urban*" OR "city" OR "cities") AND ("ecosystem service*" OR "environmental benefit*" OR "disservice*" OR "life cycle*" OR "environmental degrad*") AND ("biodivers*" OR "ecosystem divers*" OR "biological divers*" OR "species divers*" OR "ecosystem condit*") AND ( "model*" OR  "assess*" OR "tool*" OR "decision support system*"))

+ english only, article review book chapter only, 2005-2024

Scopus search

TITLE-ABS-KEY(("urban*" OR "city" OR "cities") AND ("ecosystem service*" OR "environmental benefit*" OR "disservice*" OR "life cycle*" OR "environmental degrad*") AND ("biodivers*" OR "ecosystem divers*" OR "biological divers*" OR "species divers*" OR "ecosystem condit*") AND ("model*" OR "assess*" OR "tool*" OR "decision support system*")) AND LIMIT-TO (PUBYEAR, 2024) AND (LIMIT-TO (DOCTYPE, "ar" ) OR LIMIT-TO (DOCTYPE, "re") OR LIMIT-TO (DOCTYPE, "ch")) AND (LIMIT-TO (LANGUAGE, "English"))

+ articles after 2005
