## Supplementary material for "Do urban ecosystem service assessments account for ecosystem condition and biodiversity?": SM3

### How to classify the articles

The Excel file contains the classification table in the first sheet, already filled with the 110 papers from the review, including references to the DOI links to the publications.

#### Procedure

Each row of the table initially corresponds to a paper, which may contain various models and sub-models.

To enter all available information, it will be necessary to duplicate rows, one for each method (both ES and Biodiversity), following the rules described below.

1. If the ES and Biodiversity models are **Integrated**, enter all information on a single row and answer “Yes” in the Biodiversity integration column.
   a. If the integrated model has multiple ES methods related to the same Biodiversity component, or multiple Biodiversity methods for the same ES, list the Biodiversity model(s) in a new row and answer “Yes” to **Biodiversity integration**.
2. If the Biodiversity model is not integrated, list the model(s) in a separate row and answer “No” to **Biodiversity integration**.
3. Copy all common information for each model part into each row.

#### Field Descriptions

##### Integrated

Yes/No choice

The model is integrated if: 1. The outputs of one part of the model are inputs for a subsequent part; 2. The model studies different ES but has shared equations.

##### Method type

- Biophysical
- Economical
- Socio_cultural

If unsure, check the sheets corresponding to the different method types to determine the most appropriate.

##### Method name

Dropdown list conditioned on the choice of **Method type**.

The classification of methods is available in the three sheets: Biophysical method, Economical method, and Socio_cultural method.

##### ES analyzed

- Crop provision
- Grazed biomass provision
- Livestock provision
- Aquaculture provision
- Wood provision
- Wild fish provision
- Wild animals and plants provision
- Genetic services
- Water supply
- Other provisioning services
- Carbon sequestration and storage
- Rainfall pattern regulation
- Local climate regulation
- Air filtration
- Soil quality regulation
- Soil erosion control
- Landslide mitigation
- Solid waste remediation
- Water quality regulation
- Water flow regulation
- Coastal protection
- River flood mitigation
- Storm mitigation
- Noise attenuation
- Pollination
- Pest control
- Disease control
- Habitat maintenance
- Other regulating and maintenance services
- Recreation-related services
- Visual amenity
- Education and research promotion
- Spiritual, artistic and symbolic value
- Other cultural services
- Ecosystem and species appreciation

Information on each of these ES is in the “SEEA classification” sheet.

If a known model is used (where only the calculated ES need to be specified), or if the model is unique for multiple ES, or if it is not possible to distinguish between models based on ES, more than one entry can be added by selecting the required ES.

##### Other positive or negative impact

Open-ended response.

Fill this field if the impacts (positive or negative) calculated by the model do not fall within the SEEA categories. Otherwise, leave blank.

##### Output: Quantitative, Semiquantitative or Qualitative

Select Quantitative, Semiquantitative or Qualitative according to the output provided by each method.

##### Spatial explicit

Yes/No choice.

Is the model spatially explicit?

##### Dynamical

Yes/No choice.

Is the model time dynamic?

##### Type of ES flow

- ES Potential flow
- ES Actual flow
- ES Potential & Actual flow

The ES quantity calculated is “Actual” if it includes not only what nature can potentially provide (Potential) but also what is demanded by citizens (Actual).

##### ES Sustainability Threshold

Yes/No choice.

Is there a level (or multiple levels) beyond which ES provision becomes unsustainable, i.e., overexploitation of nature?

##### Biodiversity integration

Yes/No choice.

Is the Biodiversity component properly integrated into the ES calculation? Or is it calculated separately and then compared with the quantity of ES produced to understand synergies and trade-offs?

If “No,” the two parts are essentially separate, so if multiple ES are counted, the Biodiversity part (Green) should be listed in separate rows. If integrated, it should be explained how it is integrated within the calculation, visible by the use of overlapping variables in the calculation.

##### Biodiversity level

- Genetic
- Species
- Ecosystem

At what level is biodiversity calculated?

##### Spatial scale of interest

- Local
- Regional
- Global

What scale is the biodiversity concerned with?

##### SEEA - Ecosystem condition

- A1_Chemical_State
- A2_Physical_State
- B1_Compositional_State
- B2_Structural_State
- B3_Functional_State
- _C1_Landscape

Definitions are in the “EcoCond & EBV” sheet. This selection conditions the completion of the “EBV IPBES” field.

Refers to the SEEA-EA classification where Ecosystem Conditions (EC) are defined (p. 89).

##### EBV IPBES

Dropdown menu varies depending on the previous field and is explained in the figure in the “EcoCond & EBV” sheet. All definitions are in the same sheet’s table.

Refers to Czúcz et al. (specifically in the supplementary material) comparing the SEEA-EA classification for EC and EBV. EBVs refer to type B of EC.

##### Specific biodiversity variable used

Open-ended response.

List the input variables used for calculating biodiversity. If biodiversity is integrated into the model, these should overlap at least partially with the ES model inputs.

##### Scale of model application

- Individual site
- Neighbourhood
- City
- Region (urban context at regional scale)

The scale at which the model was designed or applied.

##### NBS or ecosystem type

Open-ended response.

Specify the type of NBS the model applies to. If not applicable, specify the ecosystem type. This field is open because it is often difficult to assign the model to a single NBS. Later, responses will be analyzed and grouped into specific NBS categories if needed.

##### Case study

Yes/No choice.

Is the model applied to a case study?

##### Country

Open-ended response.

If the model is applied to a case study, specify the country. If the same model is applied in multiple countries, separate them with a comma.
